# Sparse Recovery Methods for Cell Detection and Layer Estimation

**DOI:** 10.1101/445742

**Authors:** Theodore J. LaGrow, Michael G. Moore, Judy A. Prasad, Alexis Webber, Mark A. Davenport, Eva L. Dyer

## Abstract

Robust methods for characterizing the cellular architecture (cytoarchitecture) of the brain are needed to differentiate brain areas, identify neurological diseases, and model architectural differences across species. Current methods for mapping the cytoarchitecture and, in particular, identifying laminar (layer) divisions in tissue samples require the expertise of trained neuroanatomists to manually annotate the various regions-of-interest and cells within an image. However, as neuroanatomical datasets grow in volume, manual annotations become inefficient, impractical, and risk the potential of biasing results. In this paper, we propose an automated framework for cellular detection and density estimation that enables the detection of laminar divisions within retinal and neocortical histology datasets. Our approach for layer detection uses total variation minimization to find a small number of change points in the density that signify the beginning and end of each layer. We apply these methods to micron-scale histology images from a variety of cortical areas of the mouse brain and retina, as well as synthetic datasets. Our results demonstrate the feasibility of using automation to reveal the cytoarchitecture of neurological samples in high-resolution images.

## 1 INTRODUCTION

Mapping the distribution and packing of cells (cytoarchitecture) within the nervous system is the first step in identifying the origin of tissue samples, performing comparative neuroanatomy, and delineating neurological disease states (Amunts and Zilles, 2015). Over the last century, cytoarchitectonic mappings have served as the primary way to define brain areas: researchers have estimated hundreds of cytoarchitectonically-distinct areas throughout the brain (Brodmann, 1909; Vogt, 1919; von Economo and Koskinas, 1925). These mappings have served as a standard way to perform anatomical analysis and provided neuroscientists with a standardized way of visualizing and parsing the brain, where many cytoarchitecturally distinct areas have since been correlated closely to diverse cortical functions (e.g. Brodmann areas 41 and 42 are home to primary auditory processing, Broca’s area is home to speech and language processing (Loukas et al., 2011; Ardila et al., 2016)). To this day, modeling cytoarchitecture remains an area of immense focus (Petrides, 2013; Blumensath et al., 2013; Amunts and Zilles, 2015; Weiner et al., 2017; Wagstyl et al., 2018).

A key characteristic of the cytoarchitecture of many biological tissues in mammals from mice to humans, including cerebral cortex (Belgard et al., 2011) and the retina (Wandell, 1995), is the organization of the tissues into distinct layers or “laminae”. In the neocortex there are six distinct laminae, each consisting of distinct cell types and densities of cells in each layer (Belgard et al., 2011). In the retina there are three distinct laminae: the ganglion, the inner nuclear, and the outer nuclear (Chang et al., 2007). Developing a standard framework to characterize laminar cytoarchitecture can help distinguish different brain regions and quantify disease states where the characterization of cellular layers can distinguish stages of retinal degeneration (RD) (Chang et al., 2007).

Currently, the standard method of annotating the cytoarchitecture within a given brain region relies on trained neuroanatomists to manually inscribe cells and quantify areas of interest (Gurcan et al., 2009). To characterize the laminar architecture of cortex, a study would require obtaining both the annotations of cell positions and the laminar boundaries throughout the full dataset. However, as laboratories generate larger anatomical datasets, neuroscientists require robust and efficient automated methods to facilitate the identification and modeling of cytoarchitecture.

In this paper, we present a flexible and automated framework for estimating the spatially-varying distribution of cells in a biological tissue sample directly from microscopy images. This framework consists of two main components: (i) a method for cell detection that provides robust cell detection results, even with high amounts of overlap between cells, and (ii) a method to estimate the distribution of cells that allows flexibility to find patterns in cell densities of biological importance. This framework is designed to provide a general-purpose methodology for modeling the distribution of cells in large datasets in a compact and efficient manner.

Our approach for density and layer estimation starts by modeling the presence or absence of a cell (count) as a Poisson random variable with a rate that changes according to an underlying spatially varying density function. To provide a compact representation of this potentially complex density function, we show how to represent the density as a linear combination of elements from a known basis, thus providing a generalized linear model for estimating cell densities. We then show how this modeling framework can be used to find layer boundaries (or change points) in cell densities through total-variation (TV) minimization (Rudin et al., 1992; Krahmer et al., 2017). Since the cell distributions in a close proximity of one another tend to be highly correlated, we can extend our TV-minimization approach to include a “group-sparse” penalty (Huang et al., 2011) to group together neighboring sparse estimates and therefore increase the accuracy of the estimated layer transitions in a given sample. These methods for density estimation are then brought together with a robust method for cell detection to simplify the problem of cortical layer detection to a change point estimation problem.

To test the efficacy of the proposed framework, we applied our methods to synthetic densities based upon layer estimates from somatosensory and visual cortex of the mouse/rat (Gonchar et al., 2008; Meyer et al., 2010), retinal tissue samples (Chang et al., 2007), and Nissl-stained images from the Allen Institute for Brain Science (AIBS)’s Mouse Reference Atlas (ARA) (Dong, 2008). Using the ARA, we examined the performance of our cell detection method across a variety of different brain areas, ranging from visual, auditory, motor, and somatosensory cortex, as well as retrosplenial and ectorhinal areas. Our results suggest that our methods for cell detection are comparable to the reliability across raters, and generalize well to different regions in the brain. We then test our methods for TV-minimization layer estimation on subsets of the ARA and extensively in synthetic experiments. By leveraging both a robust cell detection method for resolving overlapping cells and a state-of-the-art density estimation technique, we show that cell densities and layers can be reliably detected in high-resolution neuroanatomical images.

## 2 METHODS

Our approach for cytoarchitecture estimation, called Arcade (Approximating cellular densities), consists of three main steps (Fig. 1): (i) pulling out the cortex and surface normal directions from image data to select patches that run perpendicular to the cortical/sample surface, (ii) detecting cells in the sample and compressing these data down to spatially-distributed counts in each image patch, and (iii) estimating the density and layer transitions from the counts extracted in Steps i-ii. We provide MATLAB code and a demonstration of the framework at: github.com/nerdslab/arcade.

**Figure 1.**
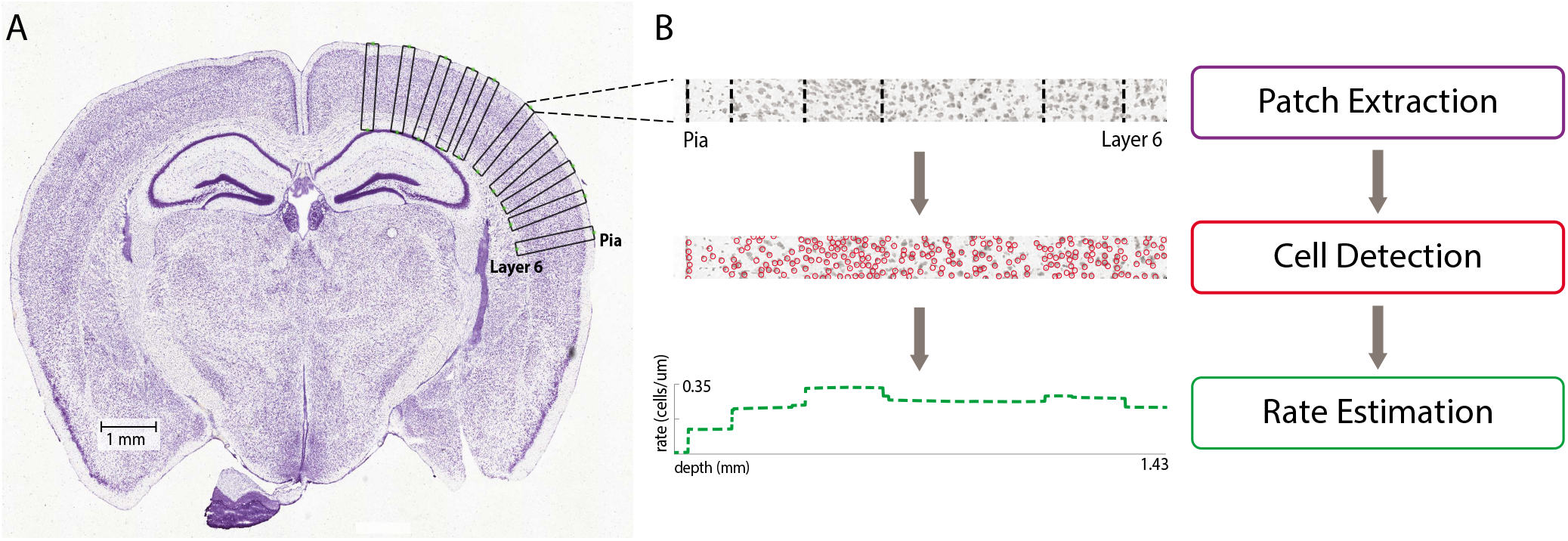
*Overview of the framework to estimate the cytoarchitectonics and layers in neuroanatomical images*. (A) A Nissl-stained image from the Allen Institute for Brain Science (AIBS)’s Mouse Reference Atlas (Dong, 2008) with a subset of detected patches overlaid around the upper right cortical surface. (B) From top to bottom: A patch extracted from the image to visualize the cellular distribution from the top (pia) to the bottom of the cortex (Layer 6). The dashed black lines indicate the transitions between layers as identified by a neuroanatomist. Below, detected cell bodies in red overlaid on the same image patch. Finally, the estimated density function obtained with a TV-minimization approach (dashed green).

### 2.1 Cortical Coordinate Extraction

The first step of Arcade involves extracting the cortex and the directions normal to the cortical surface. In the mouse brain (and other species that have a low gyrification index (Striedter et al., 2015)), sharp transitions in cell density occur when moving perpendicular to the cortical surface, noting a transition in the cellular layers (see Fig. 1). Thus, in order to estimate layers from these image data we must first determine the normal directions to the surface of the brain before layers can be estimated from a fixed coordinate system. However, due to the curved and potentially distorted surface of the neocortex, determining the direction normal to the surface proves difficult to estimate.^1^

To start, we identify the cortical/sample surface by thresholding the image and performing a connected components analysis to produce a binary image delineating the area in the image corresponding to the brain (marked ones) and the ambient background of the image (marked zeros). When calculating the surface normals, we cannot directly take the 2D gradient of the surface, as the inherent biological noise around the surface skews the approximation. Instead, we fit a smooth curve around the surface of the sample (connected components) and compute the curve’s gradient. Finally, small patches are extracted from the image that reveal the laminar architecture of the cortex and retina. In the case of the cortical sample, we split the sample into four quadrants to parallelize the patch extraction as well as to impose symmetry to mitigate edge cases for similar quarters of the sample. We use the smooth curves around each quadrant of the image to mask out the neocortex by connecting the start points and end points of each patch extracted (see Fig. 2Bi-iv), resulting in roughly 57% of the non-zero pixels of the original image to be masked. This allows us to mitigate extraneous compute-time in cell detection when we do not need information about cytoarchitecture from dense regions (e.g. the hippocampus). As a result, we extract a collection of patches from a large coronal brain section that run perpendicular to the detected and smoothed surface of the sample.

**Figure 2.**
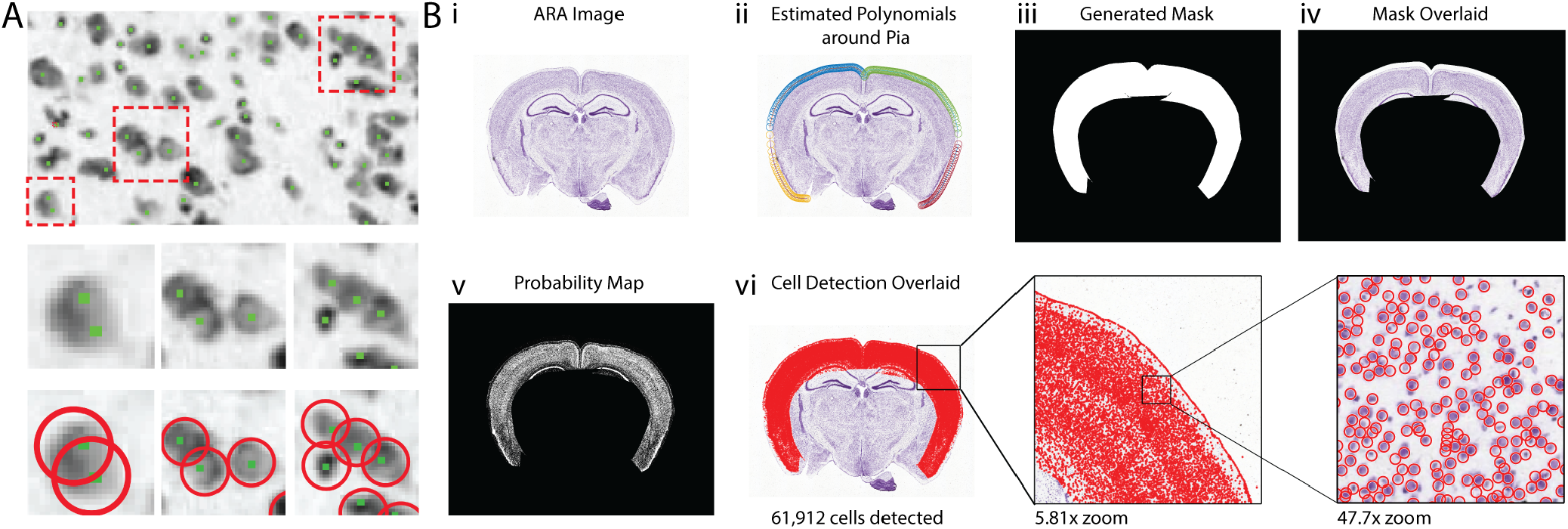
*Cortical coordinate examination and cell detection*. (A) (top row) A ground truth cutout from a neocortical sample annotated by a trained neuroanatomist at 12x magnification of a cortical slice from the ARA. The green markers indicate a cell location while the dashed red boxes show examples of overlapping cells in the ground truth. (middle row) Examples of overlapping cells at 467x, 183x, and 183x magnification of the original image. (bottom row) Estimated cells (red circles) are overlaid on the zoomed overlapping cell examples. (B) (i) Slice 293 from the ARA (0.95 *μ*m per pixel, 10,887 by 13,672 pixels). (ii) Estimated surface of the brain after fitting polynomials around the surface (pia). (iii) The mask generated to outline the cortex within the image. (iv) The mask generated in (iii) overlaid on the ARA image. (v) The probability map produced using our GMM method applied to the masked image in (iv). There are 61,912 cells detected across this section. (vi) (left) Detected cells overlaid on the original ARA image, (middle) zooming into a region at a 5.83x magnification, and (right) further magnification of a subset at 47.7x of the original ARA image (a 256 by 256 pixel cutout that spans layers 1 and 2/3 of the somatosensory cortex).

### 2.2 Cell Detection

The second step of Arcade involves processing raw histological imaging data to produce accurate cell counts and positions from the sample. Due to the thickness of the sections of the histological data in the ARA, multiple cells appear to be overlapping in the image (see Fig. 2A). For this reason, we cannot simply threshold the image to pull out all of the cells. Rather, we apply a method for greedy cell detection that can deal with overlapping cells (Algorithm 1), previously applied for for 3D X-ray neuroanatomy data (Dyer et al., 2017) and for 2D histology images (LaGrow et al., 2018) (see Fig. 2B). We now outline the two steps of the algorithm below.

To find cells in an image, we developed an unsupervised method for pixel-level segmentation that starts by converting an image into a probability map. The map encodes the probability that each pixel lies in either the foreground (cell) or background (see Fig. 2Bv). To compute a probability map, we use a Gaussian Mixture Model (GMM) to model the pixel intensities in the image. GMMs are a commonly used probabilistic model that leverage the assumption that the data in question are generated from a mixture of Gaussian distributions with unknown parameters (Reynolds, 2009). The GMM method is capable of estimating the binary pixel-level class probabilities without training data. This unsupervised method provides an efficient and accurate strategy for producing a pixel-level probability map.

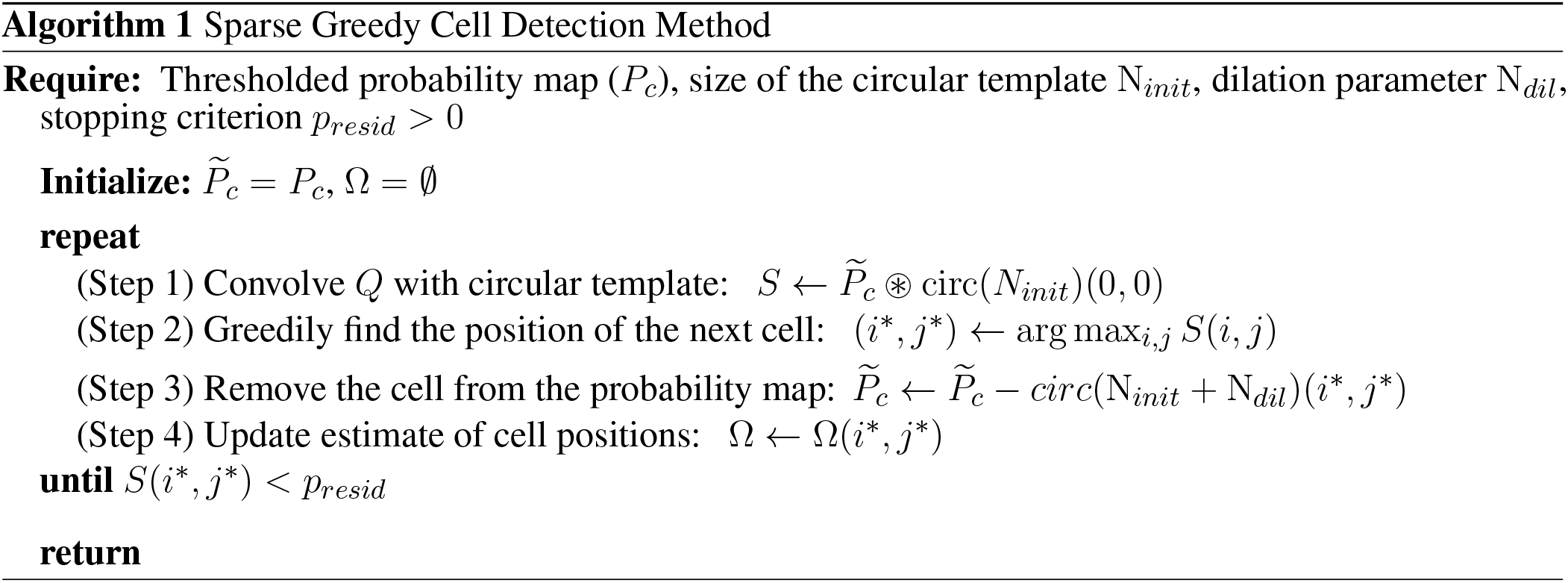

Next, we apply an iterative algorithm for cell detection designed to provide robust performance when cells are heavily overlapping (Algorithm 1) (LaGrow et al., 2018; Dyer et al., 2017). The method starts by thresholding a probability map to remove non-zero pixels with very low probability of being a cell. Then, we convolve the thresholded probability map (*P_c_*) with a circular template of fixed diameter equal to *N_init_* and find the position where the convolved map is maximum (position of next detected cell). To make this precise, the selection step to determine the spatial position of a detected cell at each iteration is given by:

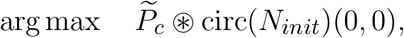

where 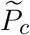 is the current probability map, circ(*N*)(*x,y*) denotes an image with a circle of diameter *N* centered at the position (*x, y*). After finding the maximum spatial position in the convolved probability map, we remove this detected cell by setting a circular region of diameter *N_init_* + *N_di1_* around the detected cell to zero, where N_di1_ is the extra amount to dilate the template to remove any residual signal. This step is performed to ensure that neighboring pixels do not influence future iterations (Fig. 2B). The algorithm continues to search for cells until a stopping criterion is reached which signifies that the correlation between the template and probability map has decreased below a certain pre-specified amount (*p_thresh_*). The output of the algorithm is a vector with the coordinates of the centers of the detected cells (see Fig. 2Bvi overlaid on the original image).

Since this cell detection method is iterative in nature and thus scales with the number of cells, high-resolution and large-scale images must be split into multiple sections and processed in parallel for more efficient computation. Once all of the sections are finished running, blocks of cell estimates are merged to produce the fully predicted map of cells.

### 2.3 Density Estimation

The third and final step of Arcade estimates a density function that matches the count data to automate the delineation of layers within a sample. A simple approach for estimating the density would entail binning up the space and counting the number of cells in each bin (Simonoff and Udina, 1997) along with smoothing methods like kernel density estimation (KDE) (Silverman, 2018). However, these histogram-based approach can be susceptible to noise and cannot exploit additional information known about the laminar organization of the tissue. Thus, we leverage the fact that the layering structure of biological tissue can be modeled by a piecewise-constant density function to find efficient representations of cellular densities.

#### 2.3.1 Density Estimation for Inhomogeneous Poisson Processes

To model the cell density in a patch of tissue as a function of the depth, we divide the space into *M* disjoint depth intervals and model the number of cells in each interval as a Poisson random variable, where *z_m_* ~ Poisson(*R_m_*) denotes the number of cells observed in the *m*^th^ bin and *R_m_* denotes the density (rate) of the *m*^th^ bin. We focus on a linear model where the density can be expressed as a linear combination of elements from a known basis ***A*** as 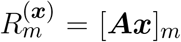, where ***A*** is a *M* × *N* matrix containing *N* basis elements each of *M* dimensions and *x* ∈ ℝ^*N*^. Under this model, the negative log-likelihood of observing ***z*** is given by

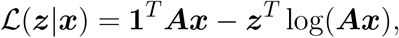

where ***z*** = [*z*_1_,…,*z_M_*] contains the number of cells in each bin and the logarithm is applied element-wise to ***Ax***.

Note that if ***A*** = ***I***, this model corresponds precisely to the simple binning approach described above, with x providing the corresponding bin counts. In general, one could choose ***A*** in a variety of complex ways to incorporate additional structure. For example, one could allow *M* to be very large (i.e., binning the space at a high resolution), but avoid the sensitivity to noise this approach would normally exhibit by keeping *N* small and choosing ***A*** to enforce a degree of smoothness across bins. Alternatively, one could design an A where N is very large, but where we can expect x to be relatively sparse, so that it can still be accurately estimated from the noisy observations.

#### 2.3.2 Maximum Likelihood Estimation with Total Variation Regularization

Estimating the density function 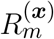 typically involves solving a MLE problem to find the parameters x that best match the observed counts ***z***. However, when dealing with noisy biological data, applying additional regularization to the estimation problem helps to mitigate overfitting.

In particular, we can model the distribution of cells across cortical and retinal samples as a piecewise-constant function, where each layer is assumed to have nearly constant density (Belgard et al., 2011). In most cases, we have strong prior information about the specific number of layers and that the number of layers *K* is small relative to the number of bins *M* used to discretize the sample, i.e., *K* ≪ *M*. Using this information, we can thus pose the density estimation problem as the following convex optimization problem:

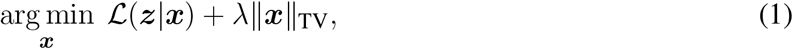

where ||***x***||_TV_ = Σ_*m*_|[***Ax***]_*m*+1_ − [***Ax***]_*m*_| is referred to as the “total variation” (TV)-norm (Rudin et al., 1992) in the basis ***A***.

When used with the TV-norm, the regularization parameter λ controls the piecewise-flatness of the resulting estimate, with larger λ will producing fewer transitions. By modulating λ, we can achieve an estimate that contains the correct number of layers in a given sample.

#### 2.3.3 Poisson TV-Sparse Algorithm

To solve the objective in (1), we employ a majorization-minimization strategy (see Algorithm 2). Majorization-minimization allows us to replace the non-differentiable total variation term with a differentiable upper bound. This allows us solve the problem via a simple iterative method, such as Newton’s method.

The majorizer we use for the total variation (in basis ***A***, denoted TV-sparse) is

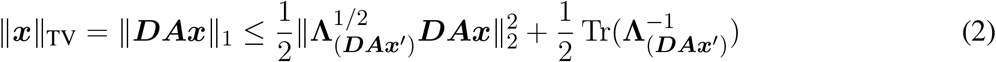

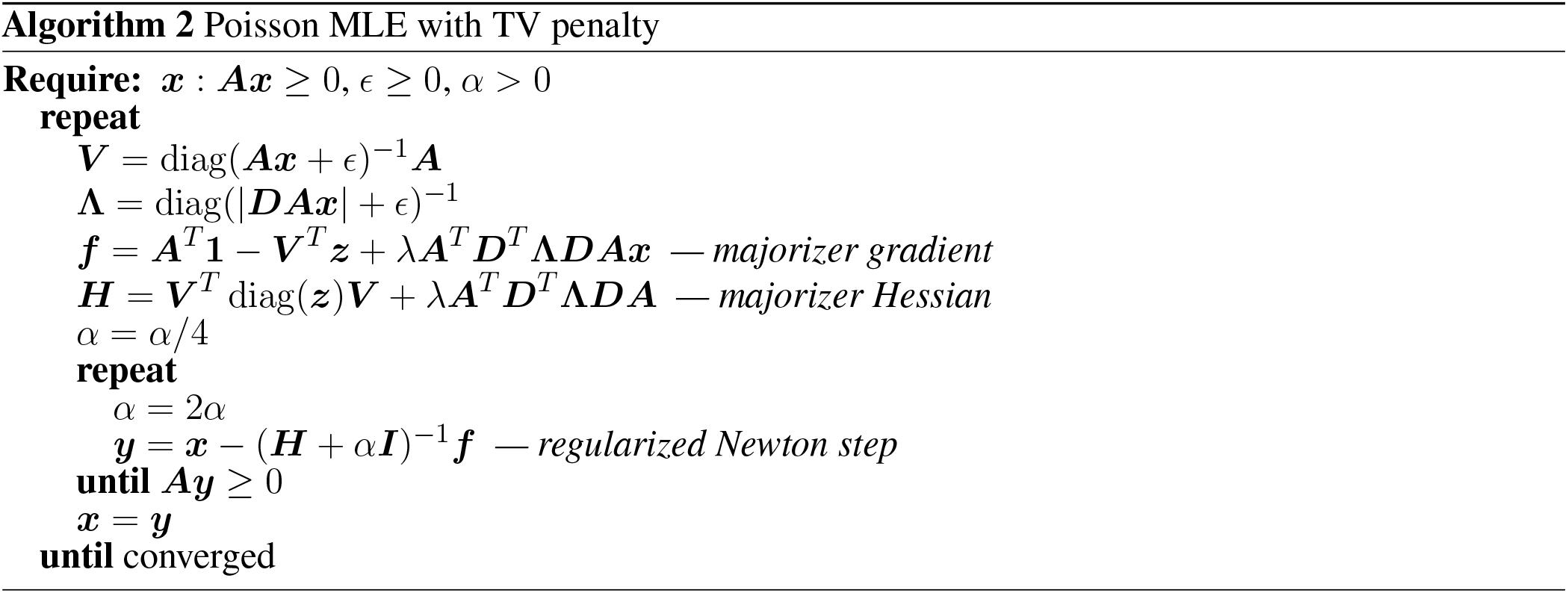

where [***DAx***]_*m*_ = [***Ax***]_*m*+1_ − [***Ax***]_*m*_ (i.e., ***D*** is (*M* − 1) × *M* and has −1 along the main diagonal and +1 along the first superdiagonal), diagonal matrix [**Λ**_(*y*)_] has entries [**Λ**_(*y*)_]_*mm*_ = |***y***_m_|^−1^, and ***x’*** is any vector of the proper size (***x*** from the previous iteration in the algorithm). The parameter *ϵ* is chosen to be: small and non-zero to condition the majorization, preventing singularities but otherwise having a negligible effect on the solution. The factor *α* is automatically tuned so that convergence is fast while accounting for the possibility of succeeding iterations estimating non-negative densities. The following reasonable initializations were used: *ϵ* = 10^−6^ · *max*(***z***) and *α* =1. Since the application uses banded matrices for both ***A*** and ***D, H*** is also a banded matrix meaning the linear system (***H*** + *α****I***)^−1^***f*** in Algorithm 2 can be solved efficiently with linear complexity of iterations in *M*.

#### 2.3.4 Group-Sparse Penalties for Spatial Regularization

The layering structure remains relatively constant within a specific neocortical region (Amunts and Zilles, I 2015). This suggests that we can potentially improve the layer estimates by using the fact that nearby patches have similar layering properties during the density estimation process. To leverage spatial similarity between nearby patches, we will need to extend the framework to jointly estimate the density of many i patches simultaneously. Suppose ***X*** and ***Z*** are obtained by stacking the patches of ***x***^(*p*)^ and ***z***^(*p*)^ into their columns, where ***x***^(*p*)^ and ***z***^(*p*)^ corresponds to the coefficients and observations of the *p*^th^ patch, respectively. The likelihood of the density of patch *p* is simply 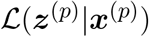. Each column (patch) within ***Z*** is independent and the total negative log-likelihood is

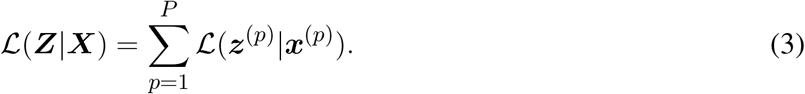

A way to extend the independent TV-minimization framework to the multi-patch case is by using the notion of group sparsity (Huang et al., 2011). With group sparsity, we can impose the assumption that layer transitions for nearby patches should occur at the same depths. This is done by sub-additively combining (e.g., via root-sum-of-squares) the variation from multiple patches (at a single depth) before summing over depths to get the total variation. Since the inter-patch consolidation is sub-additive, there is a discount for changing multiple patches at the same depth; grouped patches are encouraged to agree on layer transitions.

The penalty function we utilize is

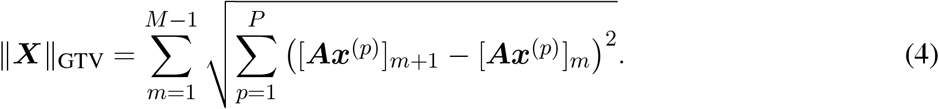

Note that, when considering *P* =1 patches, this reduces to the TV penalty discussed previously in (2).

While leveraging a group-TV penalty may appear substantially more complicated than the program outlined in (1), the resulting program can be managed by a nearly-identical algorithm by modifying a few definitions. The majorizer used for (4) is

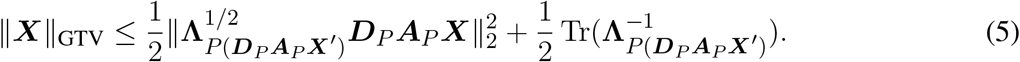

The difference matrix and basis matrix are given by the Kronecker products ***D***_*P*_ = ***I***_*P*_ ⊗ ***D*** and ***A***_*P*_ = ***I***_*P*_ ⊗ ***A***, where ***I***_*P*_ is the *P* × *P* identity matrix. Define

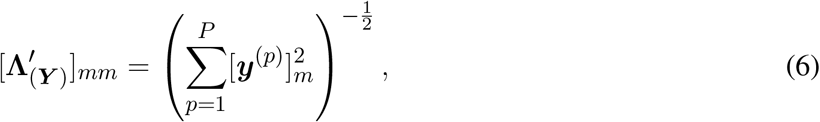

and let **Λ**_*P*(***Y***)_ = ***I***_*P*_ ⊗ **Λ’**, meaning the same coefficient is applied to [***DAx***^(*p*)^]_*m*_ at each patch *p*, but still differs by depth *m*. By replacing the corresponding substitutions to Algorithm 2 for ***Z, X, A, D*, Λ**, the resulting program will perform a group-TV-regularized Poisson maximum-likelihood estimation across multiple patches. By leveraging additional information of neighboring patches, we are able to more robustly estimate layer transitions, especially in the presence of noisy data within the image.

### 2.4 Model Selection and Parameter Optimization

To optimize our methods, we developed a hyper-parameter optimization method to ensure that the cell detection method selects the best set of parameters to find cells within the image. These parameters include: the initial threshold applied to the probability map, size of circular template (*N_init_*), size of a circular dilation window used when removing pixels corresponding to a detected cell from the probability map (*N_di1_*), and the stopping criterion for maximum correlation of the circular template within the image (P_*resid*_). This hyper-parameter optimization method uses the f1-score metric to measure the algorithm’s accuracy at correctly estimating cells given a specific combination of parameters at any given iteration. The f1-score metric is a measure based on the harmonic mean of precision (how many selected items are relevant) and recall (how many relative items are selected) (Goutte and Gaussier, 2005). The definition is as follows:

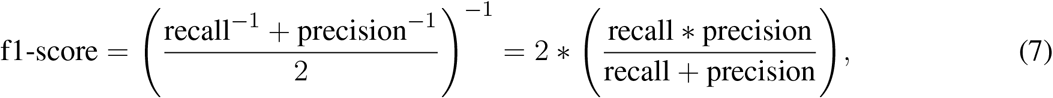

where an f1-score of 1 will be a perfect estimation of cell locations compared to the manual annotations used for ground truthing of the method. After sweeping through a range for each of the parameters, the combination of parameters that yields the highest f1-score is used to estimate cells over the entire image.

To solve the TV-minimization program to obtain a specific number of change points (layer boundaries), we: (i) fix a value of λ, (ii) run Algorithm 2, (iii) compute the number of transitions (number of non-zeros in *x*), and (iv) increase the value of λ if the number of transitions does not match the user input. We repeat this step until an appropriate λ is chosen to achieve the desired number of layers, using the solution from the previous value of λ as a warm start to the solver.

After estimating the coefficients, we often observe that transitions in density are smooth and thus produce a series or cluster of coefficients. Therefore, to indicate the transition in this case, we filter the coefficients to find local maxima and suppress the residual coefficients in a neighborhood around the maximum value. To filter the sparse coefficients, we threshold and apply a peak detector to prune the coefficients obtained through the TV-minimization method. Thereby, we are able to constrain the resulting coefficients significantly to produce an estimate with the specified number of transitions.

## 3 RESULTS

### Datasets

To test our methods on a variety of cytoarchitectonically distinct datasets, we selected four different coronal sections from the Allen Institute’s Mouse Reference Atlas (ARA) (Dong, 2008) (Fig. 3A) and an image from a retinal sample (Chang et al., 2007). The ARA is a whole brain dataset that was collected from a male C57BL/6J mouse, stained with thionin, covered with DPX, and sliced in the coronal plane. Each coronal section is 25 *μ*m thick separated by 100 *μ*m between each slice (at a resolution of 0.95 *μ*m/pixel). The second sample we tested our methods on was obtained from a retinal tissue section stained with toluidine blue (Fig. 7). The retinal dataset used in this study was obtained from a *rd10* strain of mice, an accepted animal model of retinal degeneration called retinitis pigmentosa (RP) (Chang et al., 2007). Each retinal slice is 0.5 *μ*m thick using a histo-diamond knife and heat fixed to glass slides separated by 5.0 *μ*m between each slice (at a ratio of 2.5 *μ*m/pixel). The synthetic dataset used in this paper were modeled by laminar cellular densities reported by Gonchar et al. (2008) and Meyer et al. (2010) (see Table 1). Taken together, these datasets provide a wide variety of cell sizes and packing to test our methods.

**Figure 3.**
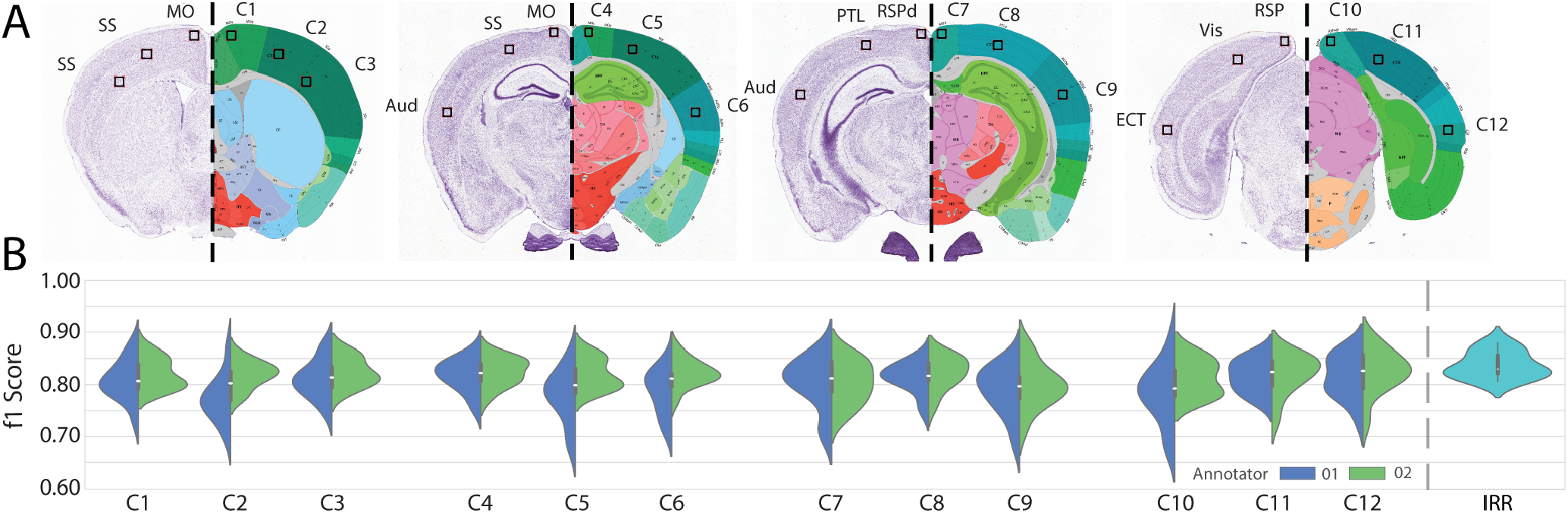
*Performance of cell detection method in different brain regions in the Allen Reference Atlas*. (A) Four coronal slices from the ARA are displayed with a dashed line mirroring half of each slice corresponding to the raw Nissl staining image and Atlas image. Each slice is inscribed and labeled with three 256 × 256 pixel cutouts marked with black squares. The corresponding brain region is marked. (B) Violin plots depicting the spread in f1-scores obtained for each cutout (C1-C12), when the automated result is compared to Annotator 1 (blue, left) and Annotator 2’s (green, right) annotations. For each cutout, we optimize and fix the hyper-parameters, and then test on all remaining 11 cutouts (that we haven’t trained on). The IRR between the annotators is displayed to the right in cyan.

**Table 1.** *Table of synthetic values for laminar transitions and cellular densities*. The values in this table were used to generate the synthetic datasets of piece-wise constant cellular layer (laminae) densities to model both the Primary Visual Cortex (V1) and the Primary Somatosensory Cortex (S1) (Gonchar et al., 2008; Meyer et al., 2010). For each cortical section, the first line indicates the normalized laminar transition starting point from the surface of the cortex (pia) while the second line indicates the normalized density of cells reported each respected layer.

| Primary Visual Cortex (V1) |  |  |  |  |  |
| --- | --- | --- | --- | --- | --- |
| Normalized Laminar Transition | 0 | 0.085 | 0.325 | 0.455 | 0.73 |
| Normalized Density of Cells | 0.042 | 0.673 | 1 | 0.56 | 0.575 |
| Primary Somatosensory Cortex (S1) |  |  |  |  |  |
| Normalized Laminar Transition | 0 | 0.079 | 0.292 | 0.45 | 0.71 |
| Normalized Density of Cells | 0.032 | 0.496 | 1 | 0.442 | 0.515 |

### Manual annotations and inter-rater reliability

After selecting four slices of interest from the ARA, we extracted six different sized cutouts (ROIs) from each coronal sections to be manually annotated. In each slice, three ROIs (512 pixels by 512 pixels) were selected and then the top left 256 pixel by 256 pixel corner of each of these larger ROIs was also selected (Fig. 3); across the four sections and six patches in each, this produced a dataset with 24 different image patches to train and test our methods. Specifically, we extracted the following image cutouts: (1) Slice 214 has one cutout from the motor cortex and two cutouts from the somatosensory cortex; (2) Slice 293 has one cutout from the motor cortex, one cutout from the somatosensory cortex, and one cutout from the auditory cortex; (3) Slice 317 has one cutout from the retrosplenial area (dorsal part), one cutout from the posterior parietal association area, and one cutout from the auditory cortex; (4) Slice 374 has one cutout from the retrosplenial area (lateral agranular part), one cutout from the visual cortex, and one cutout from the ectorhinal area. Two annotators annotated each cutout shown in Fig. 3B in ITK-Snap (Yushkevich et al., 2006) while recording the amount of time to annotate each cutout. We observe that annotations from larger cutouts have less accuracy between the two annotators, indicating additional error as the ground truth images become larger. The inter-rater reliability (IRR) (Hallgren, 2012) is comparable to the automated approach after optimization (Fig. 3B).

### Cell detection performance and generalization

Cell detection is a first critical step and thus to understand the performance of our cell detection approach across a wide range of brain areas, we utilized the many cutouts (256 x 256 pixel) described in the previous section from the ARA. We then optimized the hyperparameters used in the cell detection method to maximize the f1-score for each cutout, and tested on all other annotated cutouts (Fig. 3). Thus, the distribution of f1-scores obtained for each cutout provide insight into how well optimizing the method in each brain area will generalize to new brain areas that the method hasn’t been trained on. Additionally, we compared the IRR for the same cutouts (shown on far right in Fig. 3B) to provide an upper bound on performance within the limits of average agreement across two annotators. In most of the cutouts, we achieve similar performance with our automated approach to that of the IRR, with slightly better transfer (generalization error) for patches in somatosensory and motor cortex. Our results suggest that optimal parameters selected for single cutouts in one brain region can be transferred to new brain areas with minimal average loss in cell detection accuracy.

Additionally, we compared the timing for cell detection for both a human and our automated approach of annotation (Fig. 4). In the case of manual annotations, both annotators recorded the time that they took to annotate multiple 256×256 cutouts (left, Fig. 4A), with an average time of nearly 3 minutes. In contrast, we can run a full parameter sweep (a search over 960 different parameters) to find a set of optimal hyper-parameters for the cell detection method in approximately double the amount of time (right, Fig. 4A). Once hyper-parameters are optimized for a selected cutout, we then used the optimal set of parameters generated by the maximum f1-score to run our cell detection method on the four entire masked cortical section from the ARA (roughly 23 Million non-zero pixels, see Fig. 2). The timing of the method is then compared to the estimated amount of time needed to manually annotate the whole cortical section (12 hours, based upon linear scaling from the timing of 256 pixel by 256 pixel cutouts) (Fig. 4B). All computations were run on PC with an Intel(R) Xeon(R) CPU E3 1505M v5 (4x 2.80 GHz) and 64 GB DDR4 RAM. These results suggest that the cell detection methods can reliably detect cells from a wide variety of cortical datasets in a feasible compute-time.

**Figure 4.**
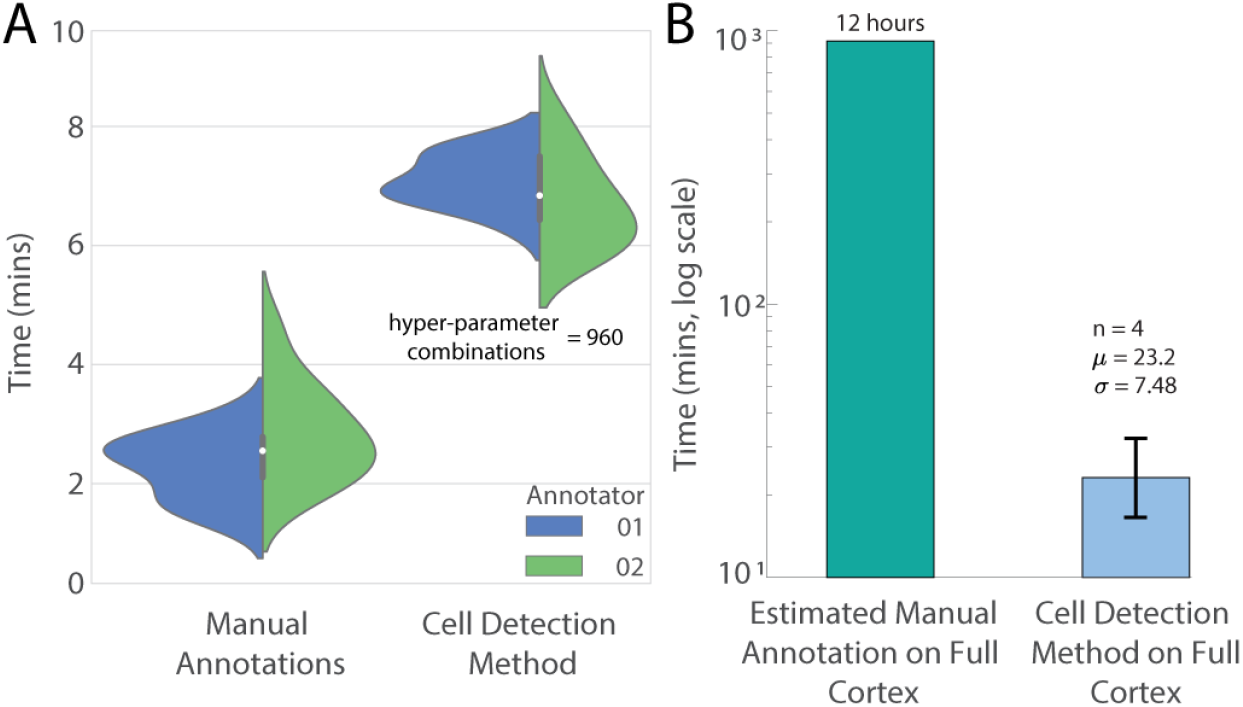
*Timing performance between manual annotations and cell detection method*. (A) Violin plots comparing the distribution of timing for manual and automated cell detection strategies. On left, manual annotations and on the right, automated annotations are displayed: for manual annotations this displays the spread in timing, for the automated result, this shows the time that it takes to run a full hyper-parameter optimization sweep (run the method 960 times to find best operating point) relative to Annotator 1’s (blue) or 2’s (green) ground truth. Each of the 960 runs of the hyper-parameter optimization takes on the order of a tenth of a second to compute. (B) The amount of time to run the method on an entire masked cortical section from the ARA (roughly 23 Million non-zero pixels). On the left, the estimated amount of time to manually annotate the whole cortical section (12 hours, based upon linear scaling from timing 256 pixel by 256 pixel patches) and on the right, the time to run the method on the same slice (average of 23 min).

### Evaluation of laminar estimation with TV-minimization

Next we tested the proposed TV-minimization approach for density and layer estimation. To do this, we initially generated two synthetic datasets that authentically modeled cellular densities in both the primary somatosensory cortex (SSp) and the primary visual cortex (VISp) in the neocortex of the rodent brain (Dong, 2008; Gonchar et al., 2008; Meyer et al., 2010) (see Fig. 5). In this case, the cortex is modeled as consisting of five layers (L1, L2/3, L4, L5, L6), each with constant density. Using this bio-realistic density function, we generated a discrete-time arrival process by drawing counts from a Poisson distribution based on either the VISp or SSp distribution function (Fig. 5). After generating counts, we then applied TV-regularization to encourage a piecewise-constant solution where the basis **A** = **I** (identity matrix), and the non-zeros in the estimated coefficients (x) correspond to the locations where each new layer begins (Table 1). Our results for both realistic cell counts in visual and somatosensory cortex show that TV-minimization can reliably estimate the density within layers and learn layer-to-layer transitions.

**Figure 5.**
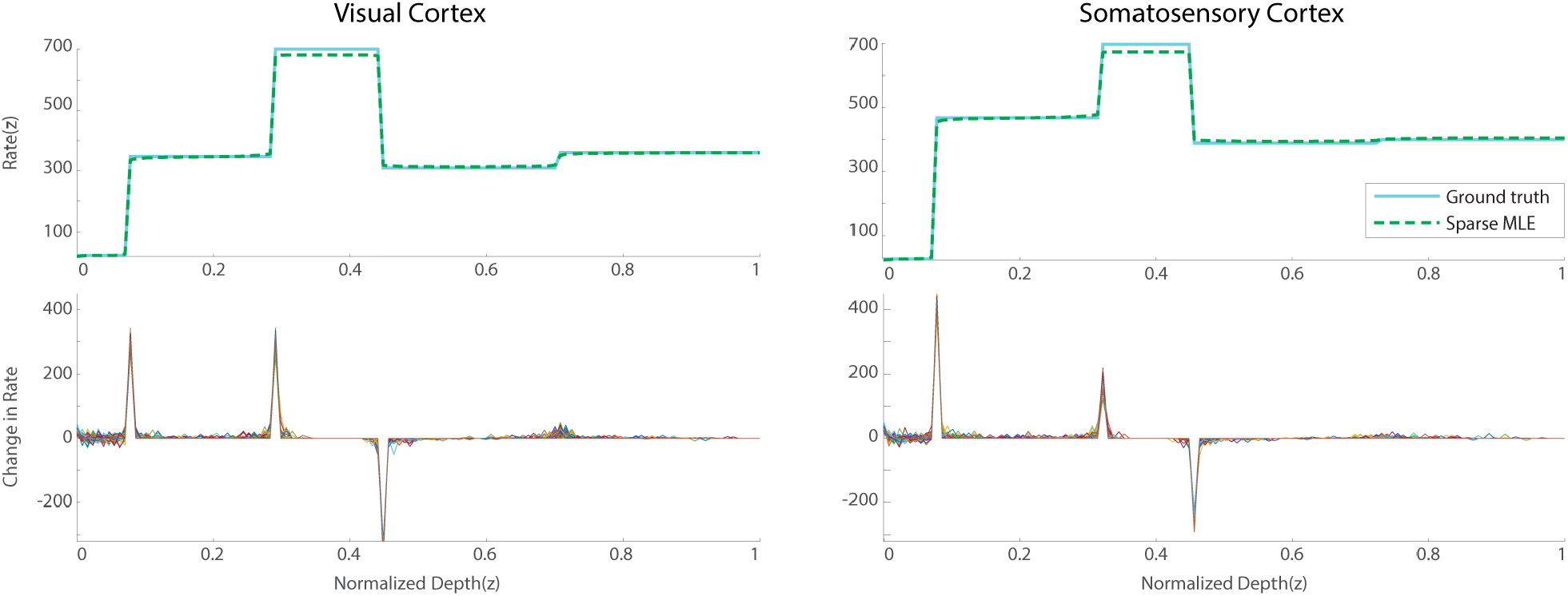
*Demonstration of the cytoarchitecture estimation framework on synthetic examples*. Synthetic cell density representations of noisy data modeled from visual cortex (left) and somatosensory cortex (right). From top to bottom: normalized density function for each cortical region (cyan) and density estimation using a sparse MLE with TV-minimization (dashed green);coefficients obtained under a sparse MLE model elucidating sparse layer transitions as measured by the total-variation norm (50 trials).

After validating that our proposed density estimation method can be applied to synthetic datasets, we applied the method to individual image patches extracted from the ARA (Fig. 6). We compared our TV-minimization approach to a more standard method for density estimation, kernel density estimation (KDE) which bins and smooths the count data (Silverman, 2018) (Fig. 6A). The accuracy of both methods was calculated by adapting a spike distance-based metric (Victor, 2005) to this setting; the spatial disparity of each transition is compared to the ground truth data by: (i) determining if the algorithm output produced the correct number of layer transitions, (ii) assigning the estimated transitions to the spatially nearest transition in the ground truth annotations, and (iii) summing the spatial pixel differences between all of the estimated and ground truth transitions. In these evaluations, both the pixel values and detected cell counts are used as input to our sparse MLE formulation for estimating the layer transitions. In the case of cell counts, the amount of cells in the same bin are added, whereas in the case of the pixel-based approach, the pixel values in a bin are added. These results confirm that utilizing TV-minimization for density estimation in conjunction with cell detection produces a more accurate estimate of layer transitions than a KDE.

**Figure 6.**
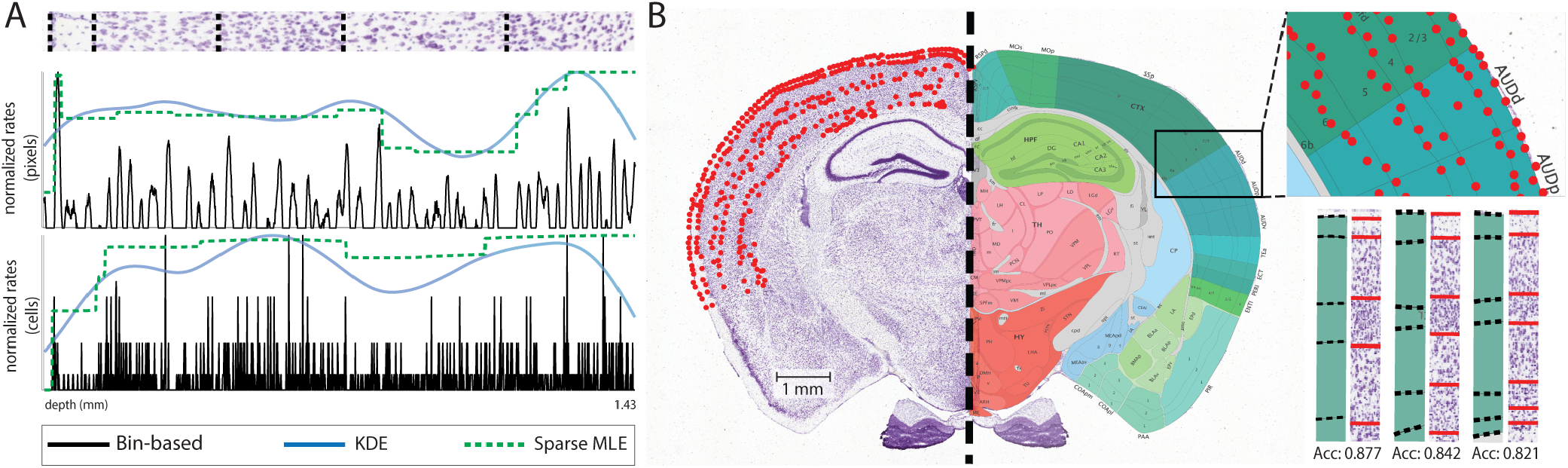
*Demonstration of density estimation*. (A) (top) An extracted patch from the primary somatosensory cortex (SSp) overlaid with the true annotated layer transitions in dashed black. (middle) Using pixel values, the plot indicate the optimal accuracy of the bin-based pixel values (black), the kernel density estimation of pixels (blue), and the sparse MLE of pixels (dashed green). (bottom) Using detected cell counts, the plot indicate the optimal accuracy of the bin-based cell counts (black), the kernel density estimation of cells (blue), and the sparse MLE of the cells (dashed green). The key corresponding to the patches and graphs follow below A. The results of layer transition accuracy are as follows for methods of bin-based, KDE, and Sparse MLE: (pixels) 0.612, 0.790, 0.822; (cells) 0.694, 0.865, 0.914. (B) The right neocortical region of the Nissl-stained image mirrored by the dashed line with both the overlaid estimated transition points in red (left) and the AIBS’s manual annotation reference atlas (right). The top inlet of B depicts the transition points overlaid against the Allen Institute Reference Atlas. The bottom inlet shows several additional patches compared to manual annotations from AIBS’s Mouse Reference Atlas using the same parameters of optimal accuracy trained in A. The accuracy of the layer transitions is displayed below each patch.

To see whether the our method of density estimation will produce a similar accuracy on new patches, we tested additional patches in the same region of the cortex as the patch from Fig. 6A (see Fig. 6B). Based on the spike-based metric of each transition compared to the the AIBS atlas (Dong, 2008), the resulting accuracy of the three additional patches were: 0.877, 0.842, and 0.821. The results demonstrate that transferring the optimal parameters from one patch to additional patches yields comparable layer transitions to the ground truth annotations provided by the ARA (see Fig. 6B). Therefore, our methods for laminar estimation are capable of providing boundaries that are comparable to the reference atlas.

### Results for retinal samples

Finally, to test our methods on a completely different type of cytoarchitecture data, we ran the proposed methods on a histopathology image of RP (retinal degeneration) from mice imaged at postnatal days 18 (P18) (Chang et al., 2007). To apply Arcade to the retinal sample, we apply the cell detection method to find cells of two different sizes since the inner nuclear layer and the ganglion cell layer of the retina have similar sized cells that differ from the size of cells in the outer nuclear layer (Chang et al., 2007). Since the staining of these images produces a slightly different range of hues for the different layers of cells (Fig. 7A), we train a three-mixture-component GMM is used to produce the probability maps of cells. The retinal sample generated an f1-score of 0.911 for the ganglion and inner nuclear layer and a f1-score of 0.900 for cells in the outer nuclear layer. After obtaining two different cell probability maps, we ran our layer estimation on the sample producing promising initial results to differentiate layer thickness and eventually characterize the degree of retinal degeneration (Fig. 7B).

**Figure 7.**
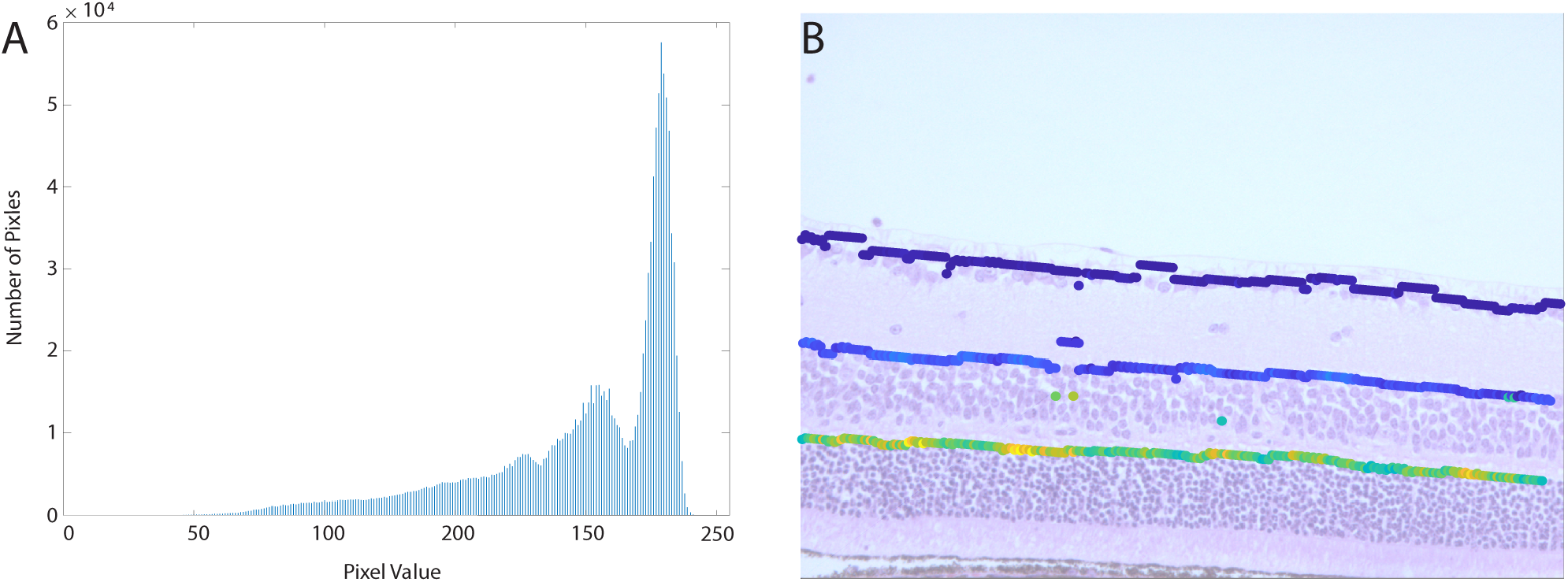
*Results of Arcade on retinal samples*. (A) Histogram of retina sample demonstrating the separability of three mixture components for the Gaussian Mixture Model. (B) Estimated layer transition points are colored according to their estimated rate (dark blue corresponds to low rates, bright yellow corresponds to high rates) overlaid on the retinal sample of a *rd10* mouse.

### Group sparsity example applied to synthetic datasets

Several artifacts and imperfections arise during slicing, sample preparation, and imaging; creating large gaps and noise in the samples (see Fig. 8A). Due to the local heterogeneity of the layer densities in the neocortex, coupling neighboring patches during layer estimation can mitigate noise presented by these artifacts. To test the hypothesis that joint estimation should improve the accuracy of density estimates over the independent case, we incorporated a group-sparse regularization penalty into the optimization framework for the constrained MLE. We tested both the independent and group-sparse methods on the VISp synthetic data set to compare both method’s performance drawn from a noisy version of the same underlying VISp density function (with Gaussian noise added to simulate potential artifacts). In the independent case, we estimated the density for each observation separately. In the group case, we estimated the density for all observations jointly. The results align well with the predictions: when we estimate the rates jointly we observe an improvement in performance (Fig. 8B) and when we increase the number of observations, we observe that the gap between the two methods also increases accordingly (Fig. 8C). These preliminary results suggest that group sparsity can be used to improve estimation of rates over the independent case.

**Figure 8.**
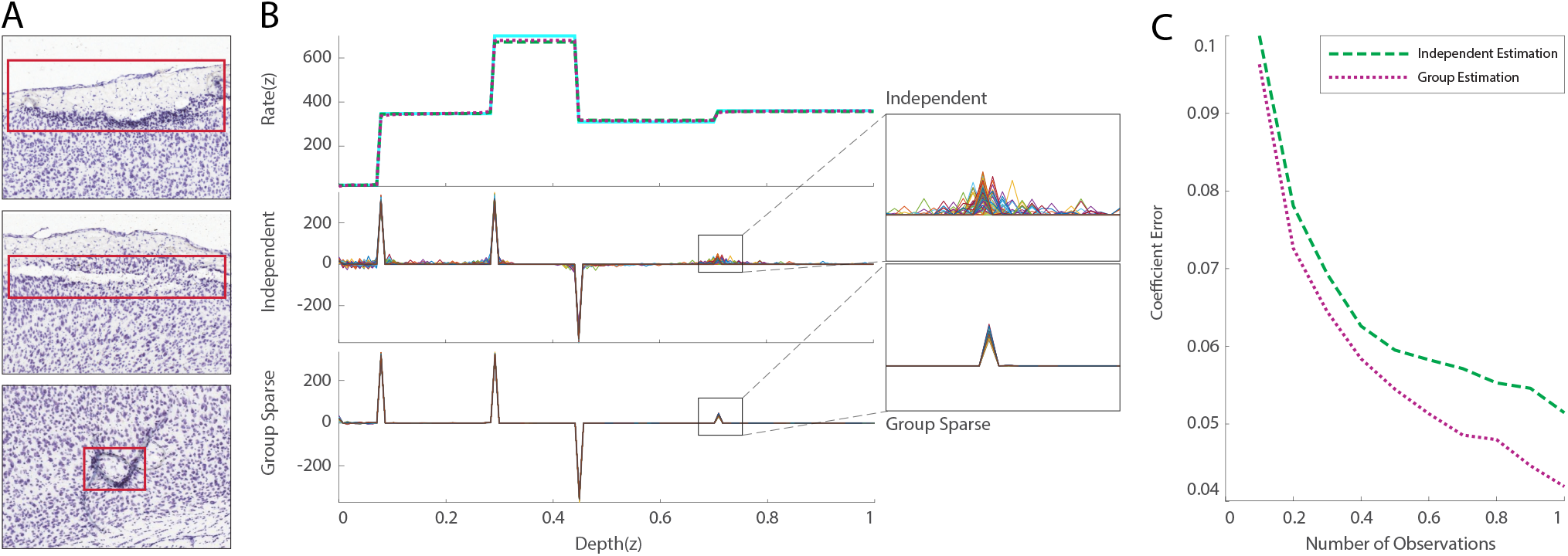
Demonstration of group-sparse approximation on three histopathology images of RP. (A) Three examples of common visual artifacts resulting from histological preparation. From top to bottom: cell death in layer 2/3; a tear in the boundary between layer 1 and 2/3; folded tissue coupled with a bubble between the coverslip and tissue. Each of these examples result in a distorted visual image of the neocortex, thereby skewing data. (B) Comparison of independent and group-sparse penalty methods. (top) The normalized density function for primary visual cortex (cyan) and density estimates obtained using an independent sparse (dashed green) and group-sparse (dotted purple) model. Coefficients obtained under an (middle) independent sparse and (bottom) group-sparse model. (C) Average coefficient error as a function of the number of observations under an independent and group-sparse model in the noisy rate setting (averaged over 15 trials).

## 4 DISCUSSION

In this paper, we introduced a framework for estimating cytoarchitecture in micron-resolution image volumes of the rodent brain. The proposed approach combines a method for cell detection with a sparse recovery-based approach for density estimation to both denoise cell density estimates and also estimate laminar transitions. We applied the framework to images from both retinal and neocortical samples and demonstrated that the methods are able to robustly detect cells and laminar transitions.

The sparse recovery framework introduced here provides a general-purpose methodology for finding structured estimates of cellular densities within various neocortical areas. We recognize the cytoarchitectonic classifications of the brain are diverse and dynamic, resulting in a number of sublayers that go beyond the 6 laminae discussed. It is of note that even within the neuroscience community, there is ongoing debate as to what is considered a “layer” within the rodent motor cortex (see Barbas and García-Cabezas (2015) for further discussion on this topic). This indicates there is a need for objective anatomical analysis which will assist anatomists in their cytoarchitectonic exploration of the brain. The framework which we have developed can efficiently and objectively educate researchers in the identification of classically defined “layers” to a high degree of accuracy.

The proposed approach for density estimation utilizes TV-regularization to find change points in cellular density. This assumes that the data consists of homogeneous regions of nearly constant density throughout the tissue. This is but one choice of a regularizer that could be used to detect layers in histological datasets. For instance, wavelets are also capable of sparsely representing density functions and promote sharp transitions in the data (Starck et al., 2010). Additionally, as we move to three dimensions, surflets (Chandrasekaran et al., 2004) could also be used to sparsely represent multidimensional density functions and promote sharp transitions in the data. By extending the method to other basis sets (beyond using TV-minimization) and to higher dimensions, the framework will increasingly be able to find transitions in densities beyond the laminar structures we considered here.

There are a number of automated methods that are tools for identifying laminae in neural datasets. These methods typically start by smoothing the image data and fitting splines or other polynomials to identify regions where the image contrast changes significantly. One such example leverages B-splines to parcellate layers in BigBrain, a high-resolution 3D model of a human brain (Lewis et al., 2014; Wagstyl et al., 2018). Similarly, in work from Feng et al. (2016), the authors developed a method to segment brain regions and layers of the olfactory bulb using a closed cubic spline (CCS)-based approach. These smoothing-based approaches have major advantages in pulling out smooth trajectories through folded cortex, however, in the process, information about specific cell counts is discarded and it thus becomes more difficult to discern small differences in the microarchitecture which can be necessary to find layers in cortex.

Recently, new tools for quantitative mapping of cell types has revealed detailed information about the morphological characteristics (i.e., shape, size, and branching structure) of cells in different cortical layers (Zeng and Sanes, 2017). The current methods do not take into account information about the shape and size of cells, however, by utilizing tools for marked point processes (Descombes and Zerubia, 2002) each of the detected counts or cells can be associated with metadata that describes different attributes of the detected cell. Thus by extending the Poisson model to a marked process model, we can capture multi-dimensional characteristics of the cytoarchitecture that includes information about cell type, shape, and size.

The methods presented in this paper could also be applied to estimate densities from a wider range of neural structures in the brain beyond cells. For instance, many efforts have been made to develop automated methods for detecting synapses (Li et al., 2010; Roncal et al., 2014; Anton-Sanchez et al., 2014; Domínguez-Alvaro et al., 2018), neuronal arbors (Sümbül et al., 2014), organelles (Perez et al., 2014), and spines (Anton-Sanchez et al., 2017). Thus, with very minor modifications sparse recovery-based methods could be used to quantify the architecture of these diverse neural structures when revealed by different imaging modalities like electron microscopy (Kasthuri et al., 2015), tissue clearing methods like CLARITY Chung and Deisseroth (2013), or X-ray microtomography (Dyer et al., 2017).

Using simulated data, we demonstrated that the proposed sparse recovery framework could be extended to the group-sparse case to jointly regularize density estimates across many nearby patches. This group sparsity approach could be further applied to help with the issue of missing data and noise (due to tears or bubbles in the tissue, common artifacts in anatomical datasets) by first detecting outlier patches and then filling in their density estimates to be consistent with their neighbors. Through utilizing other structured sparsity models (Baraniuk et al., 2010), we can extend this sparse recovery framework to take into account additional known structure like boundaries between areas.

The methods presented in this paper provide a flexible framework for automating the process of characterizing the cytoarchitecture of biological samples directly from image data. Through further developments, it will be possible to model more complex patterns of cytoarchitecture in higher dimensions and find divisions between brain areas. With automated approaches to model the architecture of the brain at multiple scales, we can further quantify changes in the cytoarchitecture due to disease (Chang et al., 2007; Hodges et al., 2006; Nobakht et al., 2011), aging (Ulrich, 1988; Roth et al., 2017), and evolution (Seelke et al., 2013).

## AUTHOR CONTRIBUTIONS

TL and ED developed the framework presented in this paper and wrote the manuscript with feedback from all authors. TL conducted the experiments and analyzed the results. MM, MD, and ED developed the methods for density estimation introduced in the paper. JP designed the synthetic datasets used in this study and annotated laminar boundaries in cortical samples. JP, AW, and TL annotated cell centers used to evaluate the performance of the cell detection method. All authors listed have made a substantial, direct and intellectual contribution to the work, and approved it for publication.

## DATA AVAILABILITY STATEMENT

The datasets and code used in this study can be found at github.com/nerdslab/arcade.

## Footnotes

1 Our current implementation relies on the fact that the sample does not contain gyri, this method could potentially be applied to human brain tissue that has been aligned to a flat map where cortex is unfolded.

